# Consensus-based somatic variant-calling method correlates *FBXW7* mutations with poor prognosis in canine B-cell lymphoma

**DOI:** 10.1101/2020.08.16.250100

**Authors:** ME White, JJ Hayward, SR Hertafeld, MG Castelhano, W Leung, SS Dave, BH Bhinder, OL Elemento, AR Boyko, KL Richards, SE Suter

## Abstract

**INTRODUCTION:** Canine Lymphoma (CL) is the most commonly diagnosed malignancy in the domestic dog, with estimates reaching 80,000 new cases per year in the United States. Understanding of genetic factors involved in development and progression of canine B-Cell Lymphoma (cBCL), the most common of the two major subtypes of CL, can help guide efforts to prevent, diagnose, and treat disease in dogs. Such findings also have implications for human Non-Hodgkin Lymphoma (NHL), as pet dogs have recently emerged as an important translational model due to the many shared histopathological, biological, and clinical characteristics between cBCL and NHL.

**OBJECTIVES:** We aimed to identify potential driver mutations in cBCL and detect associations between affected genes and differential clinical outcomes.

**METHODS:** Using exome sequencing of paired normal and tumor tissues from 71 dogs of various breeds with cBCL, we identified somatic variants with a consensus approach: keeping variants called by both MuTect2 and with high-confidence by VarScan 2. We predicted effects of these variants using SnpEff then measured associations between mutated genes and survival times from clinical data available for 62 cohort dogs using a multivariate Cox Proportional Hazards Model.

**RESULTS:** Mutations in *FBXW7*, a gene commonly mutated in both human and canine cancers including lymphoma, were associated with shorter overall survival (OS; p=0.01, HR 3.3 [1.4-7.6]). The two most frequently mutated codons of *FBXW7* in our cohort correspond to the most frequently mutated codons in human cancers.

**CONCLUSIONS:** Our findings show that exome sequencing results can be combined with clinical data to identify key mutations associated with prognosis in cBCL. These results may have implications for precision medicine in dogs and also allow subsets of dogs to serve as models for specific subtypes of human lymphoma.

**Translational Relevance:** Identifying tumor biomarkers associated with clinical outcomes has been a major driver in improved success in treating many types of human cancers, including Non-Hodgkin lymphoma (NHL). Since canine B-cell Lymphoma (cBCL) shares many clinically identifiable characteristics with NHL, our detection of recurring mutations in certain genes in cBCL and their association with clinical outcomes stands to benefit both humans and dogs. If common canine lymphoma subtypes show mutational similarity to certain human subtypes, then therapies found to be effective for a subtype in one species may be more likely to improve treatment response in the analogous subtype in the other.

## Introduction

The opportunity for pet dogs to contribute to translational research in precision medicine is unmatched by any other model species, as dogs receive the most medical assessment and intervention of any non-human animal(1). Unsurprisingly, this interest in increased medical surveillance for pet dogs comes with a demand from dog owners for continued advancements in available treatments and improved outcomes. In the case of canine lymphoma, one of the most common cancers in dogs, the parallels between canine and human disease create a unique opportunity to improve both canine and human outcomes.

Published data continues to confirm that canine lymphoma shares many clinically important characteristics with human lymphoma, including prevalence of B-cell over T-cell subtypes, existence of multiple distinguishable histologic subtypes (e.g. Diffuse large B-cell lymphoma is the most common subtype of B-cell lymphoma in both humans and dogs (2,3), varied biological behavior between cases, wide-ranging responses to treatment between individuals, and unexplained increase in incidence over the past several decades.

Biologically, pet dogs are more similar to humans than traditional mouse models in terms of size and metabolism, and this can improve the relevance of pharmacokinetic and pharmacodynamic data from drugs tested in the dog for application to human clinical research. Pet dogs also share an environment with humans and are thus exposed to similar environmental risk factors that are not present in the environments of laboratory species.

Additionally, dogs develop lymphoma spontaneously as humans do, in contrast to genetically engineered mouse models (GEMMs) predisposed to lymphoma with specific known genetic lesions or xenograft mouse models in which a human tumor is transplanted into an immunocompromised mouse (2).

Lymphoma incidence rates vary widely among dog breeds, indicating that heritable factors mediate lymphoma development. Significant progress is being made in identifying genes associated with cancer risk in dogs, providing an opportunity to better understand oncogenic mechanisms. Several genome-wide association studies (GWAS) with moderate to large cohorts of case and control dogs (Golden Retrievers [4], Standard Poodles [5], Rottweilers, and Irish Wolfhounds [6]) have revealed association signals near several known cancer genes (e.g. *TRPC6, KITLG, CDKN2A* and *CDKN2B*).

The first report of tumor mutation profiling via exome sequencing in cBCL was done using a variety of dog breeds (7). This was followed by another profiling study of exome sequencing in three breeds, the Cocker Spaniel (B-cell Lymphoma); Golden Retriever (B-cell and T-cell Lymphoma), and the Boxer (T-cell Lymphoma; 8). Recurrent mutations in B-cell lymphoma were found in *TRAF3* in both studies, validating the importance of the NF-**κ**B pathway in this disease, similar to its central role in human B-cell lymphomas.

Choosing a variant calling pipeline to identify somatic mutations in canine cancer exomes is non-trivial. First, most tumor variant calling tools are designed for use with human or rodent sequence data, so care must be taken to ensure that the algorithms can accommodate other genomes. Most striking, however, is how poorly the variant calls of multiple established callers agree with each other for the same sequence data. In a study of exome sequences from five human breast cancer patients with somatic single nucleotide variants (SNVs) and indels called by nine publicly available callers, 74% of total SNVs called (22,032 of 29,634) were called by only one caller and not detected by the other eight. Another 14% were called by only two callers. Indel calls overlapped even less with 88% (10,541 of 11,955) of indels called by only one caller and another 9% called by only two callers (9).

Gene expression profiling has likewise been applied to canine lymphoma, using a mixture of different breeds(10–13). Molecular subtypes based on dysregulated gene expression again highlighted the importance of NF-**κ**B genes, and established similarity to the human disease in the aberrant upregulation of similar pathways, although not necessarily upregulation of the same genes, as in human lymphoma(12,14). Similar to the case with mutation profiling, this has implications for biomarker development and target identification in clinical studies, and stresses the importance of developing canine-specific genomic and transcriptomic signatures.

## Methods

### Diagnosis and Immunophenotyping

Diagnosis of B-cell lymphoma was determined by board-certified veterinary oncologists and pathologists using a combination of two or more of the following methods: morphological characteristics of fixed, stained tumor biopsy; immunohistochemistry using standard anti-canine B-cell, T-cell, and other leukocyte antigen antibodies including BLA36, CD3, CD18, CD21, Pax-5, and Mac387; polymerase chain reaction (PCR) for antigen receptor rearrangement (PARR); or flow cytometry (FCM).

### Sample collection

Tissue samples from affected lymph nodes removed via excisional biopsy were provided by coauthors. Whole blood was collected at the time of excisional biopsy to be used as matched normal tissue.

### DNA Extraction

Whole blood treated with the anticoagulant ethylenediaminetetraacetic acid (EDTA) was used as the normal tissue for DNA extraction. Red blood cells were lysed, then samples were centrifuged to form a pellet of white blood cells. These cells were then lysed and treated with a salt solution to precipitate lipids, proteins, and cellular debris, leaving nucleic acids in the supernatant. DNA was precipitated in alcohol, washed to remove salt, and hydrated in a stabilizing buffer.

Formalin-fixed, paraffin-embedded (FFPE) tissue from affected lymph nodes removed via excisional biopsy were used for extraction of tumor DNA. Extraction was performed using the Qiagen QIAamp DNA FFPE Tissue Kit according to manufacturer instructions.

### Exome Library Preparation and Sequencing

Pre-capture libraries were prepared with standard library preparation protocols using the KAPA Hyper kit from Kapa Biosystems and then pooled at equal volume and sequenced on the Illumina platform at low depth to determine exact relative abundances. Based on these abundances, libraries were balanced optimally for whole exome sequencing using the SureSelect Canine All Exon V2 bait set from Agilent. Library sequencing was performed on the Illumina Hiseq 2500 platform.

### Exome Sequence Alignment

Preprocessing and alignment of exome sequences (FASTQ to BAM) were performed according to standard GATK best practices(15) using the canine reference genome file canFam3.1 (available https://github.com/auton1/dog_recomb; 16).

### Variant Calling

Several variant callers were initially tested for our dataset: VarScan 2 (17) Somatic Sniper (18), Strelka (19) and MuTect (20). A consensus approach was tested in which variants called by at least two of four callers were kept and annotated using SnpEff to predict effect. This approach, herein referred to as the 4-caller method, was evaluated via visual inspection of aligned sequencing reads at sites of variant calls using Integrated Genomics Viewer (IGV), including identification of common causes for likely false positive and false negative calls by each caller (e.g. varying minimum sequencing depth requirements, tolerance of three or more alleles at one locus, tolerance of tumor/normal contamination).

As our aim for this study was to confidently identify an initial set of common mutations that could be used to classify dogs into groups for precision treatment and ultimately match them with analogous human subgroups, we first chose to keep the conservative caller MuTect2 (21); an early, updated GATK 3-based version of the MuTect program used in the 4-caller method) because of its low frequency of false positive calls in our dataset.

Initially, when MuTect2 was run using default parameters, its relatively high frequency of false negative calls (i.e. removal of true somatic variants) was problematic. Two parameters in particular were not optimized for our samples: “max_alt_alleles_in_normal_count” and “max_alt_allele_in_normal_fraction.” Both settings aim to keep germline variants from being falsely identified as tumor variants by removing variants detected in the tumor sample that are also detected in the paired normal sample. For lymphoma, a hematopoietic cancer, tumor DNA is more likely to contaminate normal tissue (especially whole blood, the tissue used to extract normal DNA in this study) than other cancer types. The default frequency and count of sequencing reads with alternative alleles (detected in the tumor sample) tolerated in the normal sample before variants were filtered from the called variant pool were 0.03 (3%) and 1, respectively. Upon manual inspection of aligned reads in IGV, these default parameters were indeed found to be too strict; in areas of low sequencing coverage, one contaminating tumor read in the normal sample could easily represent more than 3% of total reads and in areas of high sequencing coverage, the likelihood that two or more contaminating tumor reads were present was significant. For our final variant calls, MuTect2 was run with default parameters except for the two max_alt_allele settings: max_alt_alleles_in_normal_count was changed from the default of 1 to 10,000,000 to essentially remove this filter from the variant calling process, and max_alt_allele_in_normal_fraction was changed from the default of 0.03 (3%) to 0.10 (10%). These changes decreased the frequency of false negative calls by MuTect2 in our dataset. In making these adjustments to preserve true positives, however, we also increased false positives.

In an effort to reduce false positives with minimal removal of true positives, we first used two sets of germline mutation references to remove such variants from the pool of potential driver mutations. One was a panel of normals (PON) file of germline variants generated using the exome data from normal tissue from all dogs in the study, and the other was a dbsnp file containing a catalog of germline single nucleotide polymorphisms (SNPs) called from 365 dog and wolf genomes by a coauthor’s laboratory (22). Variants that passed other thresholds to be included as tumor SNVs or indels were removed if present in the PON or dbsnp files at allele fractions greater than 10% as described above.

Next, we used a consensus approach by overlapping the remaining variants called by MuTect2 with those called with high confidence by VarScan 2. These overlaps were identified using the bcftools software package’s -isec command (23). VarScan 2 was a less conservative caller for our dataset especially in areas of high sequencing depth. Its sensitivity was an asset for our purposes to increase likelihood of finding true positive calls, while the increased false positives were likely to be filtered in the overlap with MuTect2 calls. If each were used alone, MuTect2 (with our parameter changes) and VarScan 2 called an average of 92 and 1127 variants per dog, respectively. Using the consensus method, we focused on an average of 35 mutations per dog passed by both callers.

Functional annotation of variants was performed using SnpEff(24) to process a VCF file of MuTect2 calls filtered for only those consensus mutations also called with high confidence by VarScan 2.

### Cox Proportional Hazard Model

Clinical data was gathered from medical records at the two participating referral hospital (Cornell University Hospital for Animals and North Carolina State Veterinary Hospital) and from referring veterinary records and communications where provided or available. Overall survival time (OS) was censored to the last date the patient was known to be alive in cases lost to follow up. For dogs that did not achieve remission and for dogs that were still in remission when lost to follow-up, progression-free survival time (PFS) was set to missing and not included in calculations.

First, a univariate Cox regression was performed with the clinical data available from 62 of the dogs in the study using covariates (other than mutation status) known to affect survival times in cancer patients (e.g. age, sex, initial treatment). The Cox proportional hazards model used to test OS with mutation status by gene was fitted with non-colinear covariates that had demonstrated effects in the univariate models, including age at diagnosis, sex, body weight at time of diagnosis, and initial treatment factor (no treatment, induction chemotherapy only, full course of single-agent chemotherapy, or full course of multi-agent chemotherapy). The Cox proportional hazards model used to test PFS with mutation status by gene was fitted with the same covariates plus another indicating treatment status at relapse (treated or untreated). A mutation in a gene was considered to have a significant effect on time from remission to relapse (PFS) or time from diagnosis to end of life (OS) if the p value of the mutation status’s coefficient in the model was less than 0.05. Data was input into the R packages “survival”(25) and “survminer” (26).

## Results

Compared to our initial 4-caller method (Supplemental Figure 1, Supplemental results), our final method using consensus variants called by both MuTect2 and with high confidence by VarScan 2 yielded many fewer variants, but kept variants were much more likely to be true positive somatic mutations when sequencing read alignments were visually inspected.

In our dataset, *TRAF3* was the most commonly mutated gene in cBCL (n=20, 28% of dogs). This is similar to a previous study sequencing cBCL in the Golden Retriever and Cocker Spaniel in which 13 dogs of 64 (20.3%) had at least one somatic mutation in *TRAF3* (8). TRAF3 has a diverse role as a regulator of both the canonical and non-canonical NF-κB pathways known to play a role in the pathogenesis of many cancers(7,27). In our dataset, *TRAF3* mutations were not associated with significant differences in overall survival or progression-free survival.

**Table 1:**
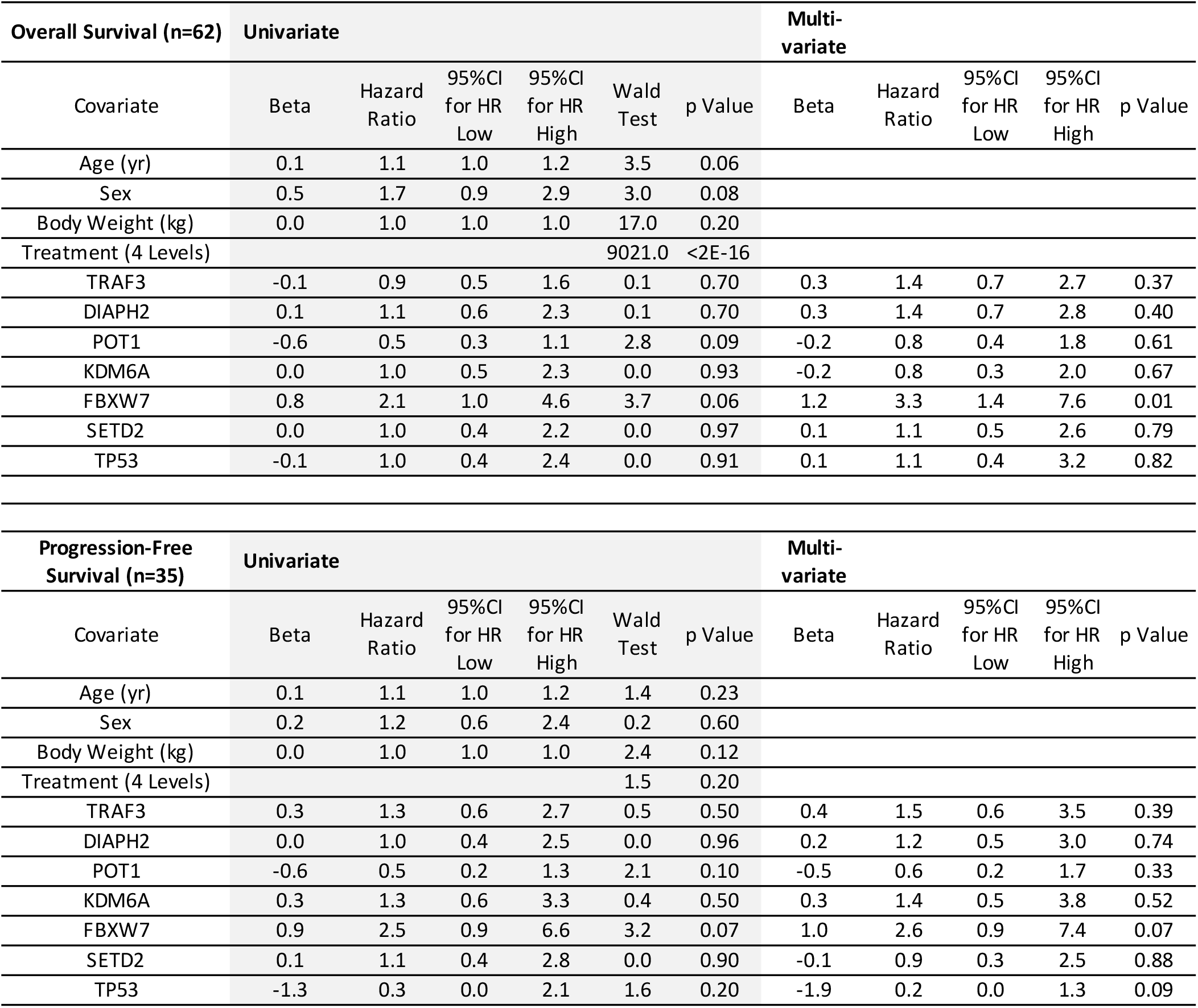
Results of Univariate Cox Regression and Multivariate Cox Proportional Hazards Model

The next most frequently mutated gene was DIAPH2 in 17% of dogs (n=12), with only 2 dogs having missense variants with moderate predicted effect and the rest of dogs having one or more mutations in introns of the gene. In dogs with clinical data unaffected by experimental treatments or additional types of cancer (n=62), mutations in DIAPH2 were not associated with significant changes in OS or PFS.

*ENSCAFG00000030258*, part of the immunoglobulin heavy chain, mu (IGHM) was the next most frequently mutated gene with all mutations synonymous or outside of coding regions in 15% of dogs (n=11). These mutations are not likely to be drivers in the transition to malignancy and may instead be due to either hypermutation in the germinal center or allelic variation(3). This is also likely true for another highly mutated gene, *ENSCAFG00000029236*, part of the immunoglobulin light chain, lambda (IGL) with all mutations synonymous or outside of coding regions, found in 14% of dogs (n=10).

*POT1* (protection of telomeres 1), a component of the telomerase ribonucleoprotein (RNP) complex necessary for proper replication of chromosome ends, was mutated in 14% of dogs (n=10). *POT1* has previously been implicated in cBCL, with 11/64 (17%) dogs with BCL having non-silent, protein-coding mutations in one study of cBCL in two dog breeds (8). All 10 dogs with *POT1* mutations in our study had a single mutation in the gene with moderate to high predicted effect. However, for dogs in our study, *POT1* mutations were not associated with significant differences in overall survival or progression-free survival.

*KDM6A* (lysine demethylase 6A) was mutated in 13% of dogs (n=9). Eight of these mutations were in coding regions with moderate (n=3) or high (n=5) predicted effect and one dog (ID 4101) had a splice region and intron variant with low predicted effect. The five mutations with high predicted effect were annotated as loss-of-function (LOF) variants, which is of translational significance given that gain-of-function mutations in *EZH2* (Enhancer of zeste homolog 2), a gene encoding a histone-lysine N-methyltransferase enzyme, have been identified in approximately 22% of diffuse large B-cell lymphoma (DLBCL) cases in humans. (28) EZH2 has an opposing effect to KDM6A, and therefore the presence of LOF mutations in *KDM6A* may make these dogs an appropriate translational model for human cancers with GOF mutations in *EZH2*. Pairing with clinical data did not show that KDM6A mutations associated with significant differences in overall survival or progression-free survival.

*FBXW7*, F-box and WD repeat domain containing 7, was mutated in 13% of dogs (n=9 dogs), with 2 dogs each having 2 separate mutations (11 total mutations). The first of these two latter dogs (ID 4117) had one frameshift mutation in codon 231 of 712 with high predicted effect and one missense mutation at amino acid 484 of 712 with moderate predicted effect. Canine codon R484 corresponds to one of the two most commonly mutated human codons of *FBXW7* in cancer, codon R479 (29). The second dog with two mutations in *FBXW7* (ID 4127) had one premature stop codon at amino acid 229 of 712 with high predicted effect and a frameshift mutation at amino acid 329 of 712 with high predicted effect. Both dogs were heterozygous for each of their two mutations with no fragments spanning both variant sites, so it is not known for either dog whether the dog has two mutations in one copy of the *FBXW7* gene (and one wildtype copy) or whether the dog has two mutated copies of *FBXW7*. For the 7 dogs with a single mutation in *FBXW7*, 6 had mutations in coding regions with moderate or high predicted effect, and one dog had a mutation 9 base pairs upstream of the transcription start site (i.e. in the promoter region).

Dogs with mutations in *FBXW7* had significantly shorter overall survival using data from 62 dogs with OS data (p=0.01, Hazard Ratio 3.3; Figure 3) and reduction in progression-free survival (PFS) that approached significance in the small subset of 35 dogs with known achievement of remission and known relapse dates (p=0.07. Hazard Ratio 2.6; Figure 3).

**Figure 1:**
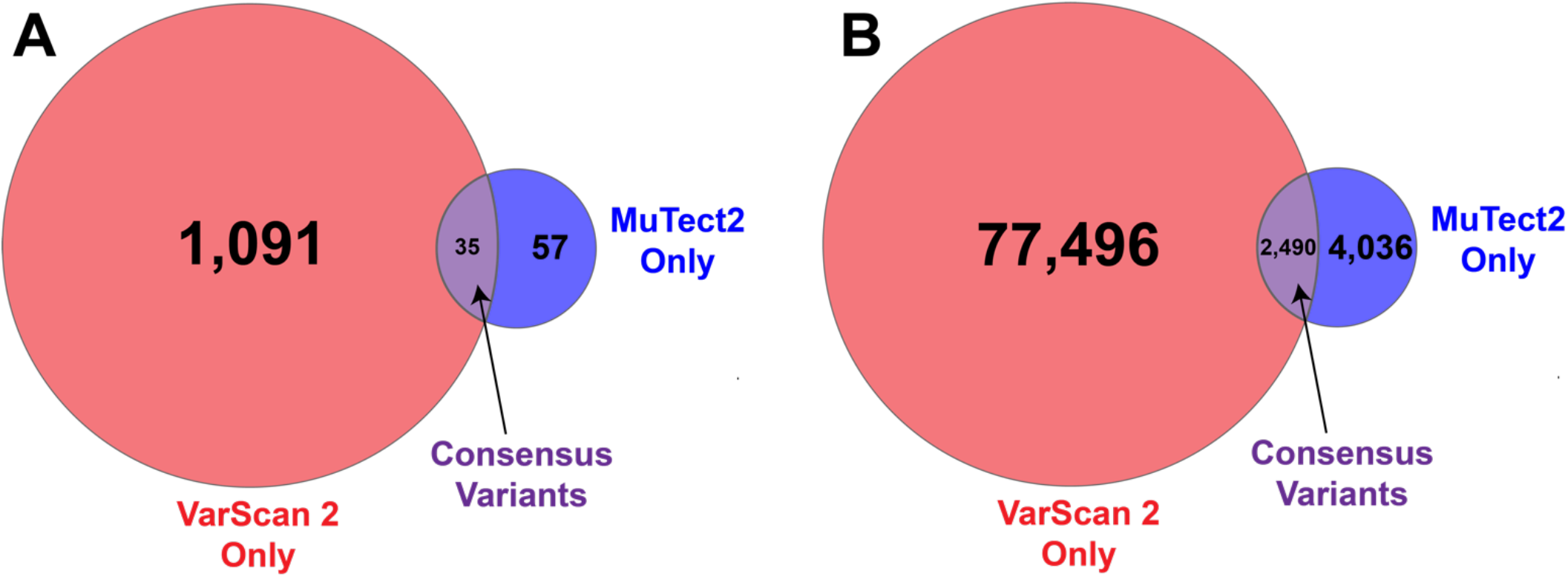
Number of tumor variants kept by individual callers and using consensus method shown as (A) average per dog and (B) combined for all dogs (n=71). Consensus variants shown in purple. Variants called by only VarScan 2 with high confidence in red and variants called by only MuTect2 with adjusted parameters (see methods) in blue.

**Figure 2:**
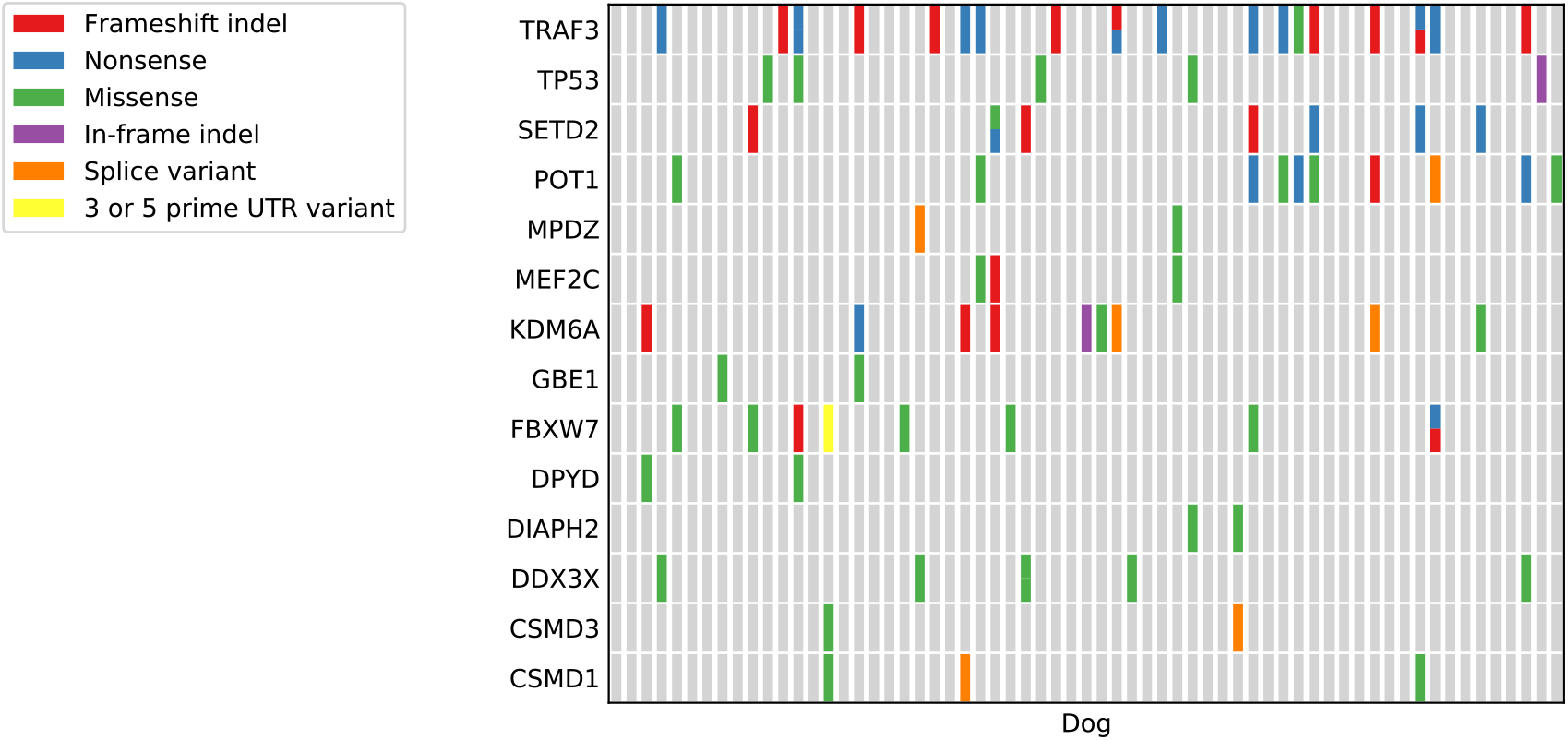
Map of selected functional categories of somatic mutations in most recurrently mutated genes. Each column represents an individual dog, and colored bars indicate the dog had one or more somatic mutations called via 2-caller consensus method in that gene. Only mutations with greatest predicted effect are shown.

**Figure 3:**
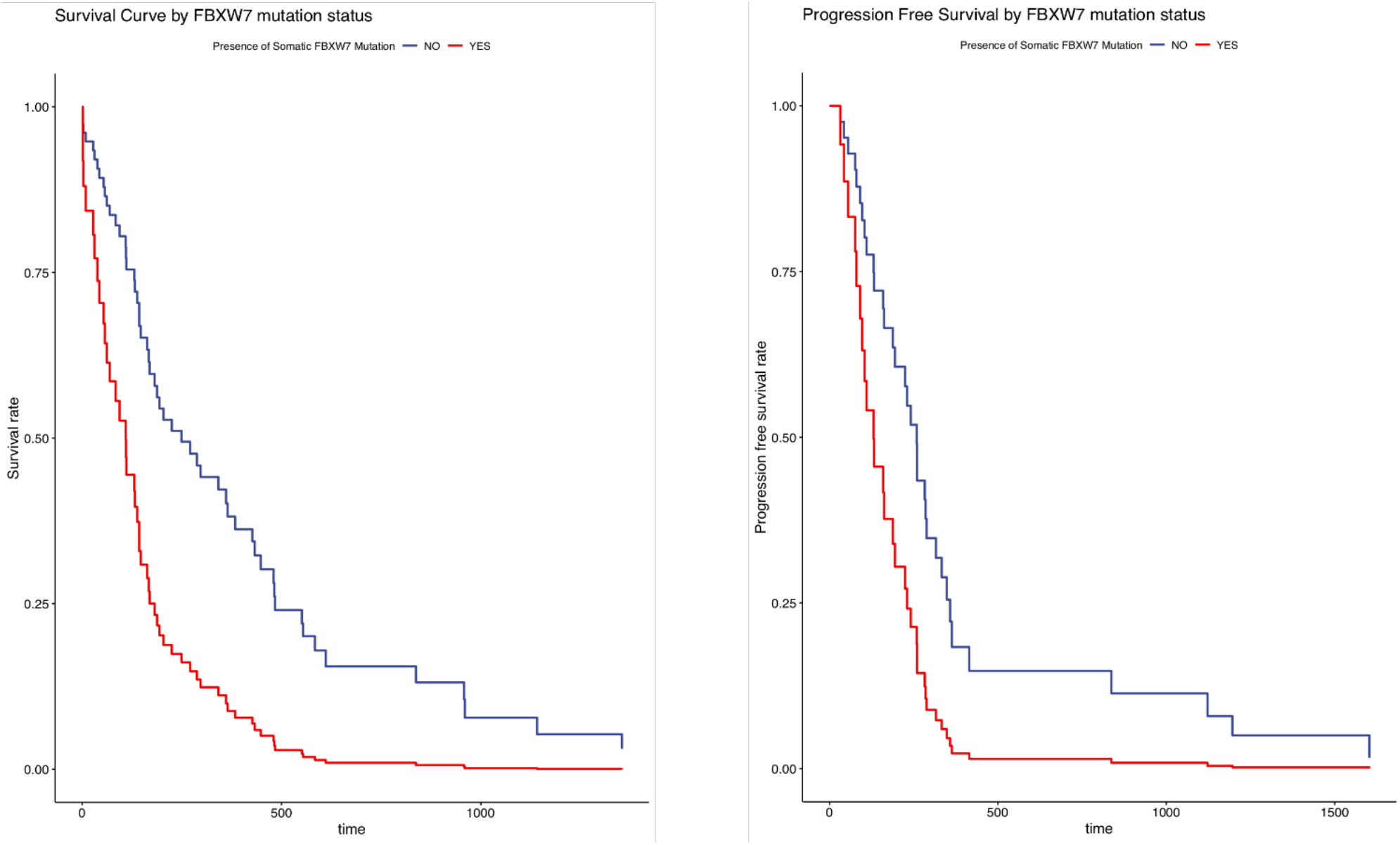
Survival curve (left, n=62) and Progression-free survival curve (right, n=35) of dogs with (red) or without (blue) a somatic mutation called in *FBXW7*. Dogs with a called somatic mutation in *FBXW7* had a significantly shorter overall survival time (p=0.006, HR 3.26) and a shorter progression-free interval that approached significance (p=0.07).

In a previous study of canine lymphoma from three dog breeds (8) *FBXW7* was the second most commonly mutated gene in canine B-cell tumors of the Golden Retriever and Cocker Spaniel combined and most commonly mutated gene in cBCL tumors of the Golden Retriever alone. Forty-one percent of these mutations were in canine codon R470. This codon in the dog corresponds to the second of the aforementioned two most commonly mutated human codons, R465 (29). In our study, 2 dogs had tumors with missense mutations in canine codon R470, and 2 (as mentioned above) had mutations in the other corresponding commonly mutated codon, but the remaining 7 mutations were elsewhere. Four other mutations were also in the sequence encoding the WD40 repeating domain of the FBXW7 protein (both of the two mutations in dog 4127 and one mutation in dog 4117 described above plus one dog with a frameshift at canine codon 401), and two dogs both had a missense mutation in canine codon 524.

*FBXW7* is commonly mutated in many human cancers (mutation frequency across all types has been reported at approximately 6%, with variation by cancer type; 30). Mutations are frequently found in a heterozygous state and, because of FBXW7’s action as a dimer, these mutations can still easily affect protein function in terms of stability and substrate affinity (31). Mutations in *FBXW7* reducing protein function have been implicated in resistance to chemotherapy and poor prognosis in human cancers including non-small cell lung cancer and pancreatic cancer due to stabilization of antiapoptotic protein MCL-1 (32,33).

*SETD2* was mutated in 11% of dogs (n=8). One dog (ID 4167) had 2 mutations in *SETD2*, one with a premature stop codon at amino acid 1541 of 2562 with high predicted effect and the other a missense variant with moderate predicted effect at amino acid 1578. The dog is heterozygous for both mutations, and it is unknown whether the dog has two mutations in one allele of *SETD2* (and one wildtype copy) or whether the dog has two mutant *SETD2* alleles. For the 7 dogs with a single mutation in *SETD2*, all had mutations in coding regions with high predicted effect. SETD2 has many diverse roles in the body and thus mutations can have a variety of phenotypes including loss of genomic stability, disruption of the p53 apoptosis response, changes to recruitment of splice machinery and splice sites used, and altered recruitment of proteins in DNA damage response (34). Mutations in *SETD2* were not significantly correlated with overall survival or progression-free survival in our study.

*TP53* was mutated in 8% of dogs (n=6). This tumor suppressor is the most frequently mutated gene detected in human cancers with over half containing a mutation in the gene (35,36), and it has also been shown to be frequently mutated in canine cancers including lymphoma (1), for which the mutations have been associated with decreased response to treatment and shorter survival times(37). In our cohort, *TP53* mutation was not associated with significant changes to overall survival time, but a decrease in the progression-free survival for dogs with a mutation in the gene approached significance (p=0.09). Due to its common presence in many diverse human cancers, TP53 function is repeatedly targeted for cancer therapies. The common presence of *TP53* mutations in dogs could make them more useful for development of such targeted therapies.

## Discussion

In our initial attempts to optimize a variant calling pipeline for exome sequencing of canine lymphoma, we used a consensus approach of accepting variants called by 2 or more of 4 variant callers. Overlap between callers was very small and even kept calls had a high likelihood of being false positives when manually curated. We adjusted our approach using MuTect2, an updated version of the caller that had performed best in our previous attempt in terms of calling mostly true positive calls as determined by visual inspection. However, we found that this caller often rejected true positive mutations due to tumor contamination in the normal sample reads, which the caller interpreted as the mutation being germline in origin. As tumor contamination into normal tissue is a common problem in hematopoietic cancers(38), we first relaxed standards for rejecting mutations found in normal tissue as germline mutations to allow for a small amount of contamination of tumor tissue into normal tissue. We reduced the introduction of false positives created by this change by only accepting mutations scored as high confidence by VarScan 2, a less conservative caller that performed well in our initial approach with a greater likelihood of keeping true positives compared to MuTect but simultaneous increase in the calling of false positives. By using the extremely conservative consensus approach of these two callers, we were still able to identify commonly mutated genes in canine B-cell lymphoma in our dataset and found a significant relationship to clinical outcomes based on mutations in one gene. This would be expected if the set of mutations identified in our study was enriched for actual “driver” mutations important to the establishment and progression of lymphoma (rather than “passenger” mutations arising due to genome instability or other malignancy factors that have little to no effect on pathogenesis (39) that confer a selective advantage on that cell and its offspring. These cells would become the predominant population and thus their mutations will be sequenced more frequently and called with increased frequency and confidence by variant callers.

Dogs with mutations in *FBXW7* show promise for translational research because of their similarity to humans in terms of codons affected and the shared, negative effect of coding mutations on prognosis. Prevalence of *FBXW7* mutations differed by breed in a study comparing exome sequencing data with Golden Retrievers much more likely to have a mutated allele than Cocker Spaniels with B-cell lymphoma (8). No two dogs of the nine with *FBXW7* mutations in our study were from the same breed (though one was a purebred Labrador Retriever and one was a Labrador Retriever cross) but our sample size was small. Continued profiling of canine lymphoma is necessary to reveal if breed may be useful as a proxy for tumor profile in making clinical decisions in canine lymphoma.

Dogs with mutations in *TP53* are also poised to contribute to the development and testing of drugs. Because of its overwhelming frequency of mutations in human cancers, intense focus has been placed on developing new drugs targeting *TP53*-mutated cancers. With its diverse potential contributions to malignancy, many options exist for potential mechanistic targets for changes to TP53 activity. With so many resources aimed at such drugs and so many affected dog and human patients, it is crucial to test each potential therapy for safety and efficacy as soon and as quickly as possible. Here, the condensed lifespan of the dog compared to human could be of great benefit, as tumors occur and dogs relapse over a shorter span of time such that studies could be completed in less time (2).

Future directions for this work should include the study of differential gene expression by somatic mutation to increase understanding of the downstream effects of these variants and the biological pathways involved in tumor progression. Additionally, gene expression work in cell lines with single gene knockouts or GOF mutations in human and dog cell lines will provide a more controlled experiment regarding the effects of each somatic mutation.

Information from this study increases our potential to create canine lymphoma subtypes based on genes mutated, and this will aid in assessing their similarity to human lymphoma subtypes. Additional clinical genomic data is needed with dogs phenotyped by breed, histopathological lymphoma subtype, and clinical outcomes. Such data will also help establish whether differences between our study and others in terms of codons affected, differences in the frequency that specific genes are mutated, and effect of mutations on clinical outcomes is due to differences in breeds sampled or the heterogeneity of the canine lymphoma mutation landscape.

The outlook for dogs becoming increasingly valuable as translational models for lymphoma continues to improve as dogs receive increased medical surveillance and treatment, costs of sequencing decrease, and additional molecular similarities between human and canine lymphoma are discovered. Studies like the Golden Retriever Lifetime Study(40) may help with the contribution of serial biological samples (metabolites, cfDNA, circulating tumor cells) that can be used to develop and improve more sensitive detection methods for the development and/or return of cancer. Additionally, such studies could bank healthy tissue earlier before cancer develops in the bloodstream, thus preventing many of the issues with tumor contamination in normal tissue leading to rejection of true somatic mutations as germline variants.

As clinical data from more dogs is paired with genomic data, we will be able to begin making precision recommendations for dogs with certain mutation profiles. As these mutations are further researched and studied, we will gain a better understanding of how these mutations affect gene expression and tumor behavior and gain insight into the mechanisms of cancer pathogenesis and differential response to treatments between types. Ultimately, this work can improve both canine and human outcomes in patients with lymphoma.

## Supporting information

Supplemental Results

## Acknowledgments

The authors thank the dogs and owners that participated in this study. We also thank Susan Garrison and the Medical Genetics Service at Cornell University Hospital for Animals (CUHA) for their assistance with tumor and record collection for many dogs in this study. We appreciate Dr. Michael Boyle for his assistance with python scripts.

## References

1. Ostrander EA, Dreger DL, Evans JM. Canine Cancer Genomics: Lessons for Canine and Human Health. Annu Rev Anim Biosci. 2019;7:449–72.

2. Richards KL, Suter SE. Man’s best friend: what can pet dogs teach us about non-Hodgkin’s lymphoma? Immunol Rev. 2015;263:173–91.

3. Avery AC. The Genetic and Molecular Basis for Canine Models of Human Leukemia and Lymphoma. Frontiers Oncol. 2020;10:23.

4. Hayward JJ, Castelhano MG, Oliveira KC, Corey E, Balkman C, Baxter TL, et al. Complex disease and phenotype mapping in the domestic dog. Nat Commun. 2016;7:10460.

5. Karyadi DM, Karlins E, Decker B, vonHoldt BM, Carpintero-Ramirez G, Parker HG, et al. A Copy Number Variant at the KITLG Locus Likely Confers Risk for Canine Squamous Cell Carcinoma of the Digit. Plos Genet. 2013;9:e1003409.

6. Karlsson EK, Sigurdsson S, Ivansson E, Thomas R, Elvers I, Wright J, et al. Genome-wide analyses implicate 33 loci in heritable dog osteosarcoma, including regulatory variants near CDKN2A/B. Genome Biol. 2013;14:R132.

7. Bushell KR, Kim Y, Chan FC, Ben-Neriah S, Jenks A, Alcaide M, et al. Genetic inactivation of TRAF3 in canine and human B-cell lymphoma. Blood. 2014;125:999–1005.

8. Elvers I, Turner-Maier J, Swofford R, Koltookian M, Johnson J, Stewart C, et al. Exome sequencing of lymphomas from three dog breeds reveals somatic mutation patterns reflecting genetic background. Genome Res. 2015;25:1634–45.

9. Krøigård AB, Thomassen M, Lænkholm A-V, Kruse TA, Larsen MJ. Evaluation of Nine Somatic Variant Callers for Detection of Somatic Mutations in Exome and Targeted Deep Sequencing Data. Plos One. 2016;11:e0151664.

10. Frantz AM, Sarver AL, Ito D, Phang TL, Karimpour-Fard A, Scott MC, et al. Molecular Profiling Reveals Prognostically Significant Subtypes of Canine Lymphoma. Vet Pathol. 2013;50:693–703.

11. Mooney M, Bond J, Monks N, Eugster E, Cherba D, Berlinski P, et al. Comparative RNA-Seq and Microarray Analysis of Gene Expression Changes in B-Cell Lymphomas of Canis familiaris. Plos One. 2013;8:e61088.

12. Mudaliar MAV, Haggart RD, Miele G, Sellar G, Tan KAL, Goodlad JR, et al. Comparative Gene Expression Profiling Identifies Common Molecular Signatures of NF-κB Activation in Canine and Human Diffuse Large B Cell Lymphoma (DLBCL). Plos One. 2013;8:e72591.

13. Su Y, Nielsen D, Zhu L, Richards K, Suter S, Breen M, et al. Gene selection and cancer type classification of diffuse large-B-cell lymphoma using a bivariate mixture model for two-species data. Hum Genomics. 2013;7:2.

14. Richards KL, Motsinger-Reif AA, Chen H-W, Fedoriw Y, Fan C, Nielsen DM, et al. Gene Profiling of Canine B-Cell Lymphoma Reveals Germinal Center and Postgerminal Center Subtypes with Different Survival Times, Modeling Human DLBCL. Cancer Res. 2013;73:5029–39.

15. Auwera GA, Carneiro MO, Hartl C, Poplin R, Angel G del, Levy-Moonshine A, et al. From FastQ Data to High-Confidence Variant Calls: The Genome Analysis Toolkit Best Practices Pipeline. Current Protocols in Bioinformatics. 2013;43:11.10.1–11.10.33.

16. Auton A, Li YR, Kidd J, Oliveira K, Nadel J, Holloway JK, et al. Genetic Recombination Is Targeted towards Gene Promoter Regions in Dogs. Plos Genet. 2013;9:e1003984.

17. Koboldt DC, Zhang Q, Larson DE, Shen D, McLellan MD, Lin L, et al. VarScan 2: Somatic mutation and copy number alteration discovery in cancer by exome sequencing. Genome Res. 2012;22:568–76.

18. Larson DE, Harris CC, Chen K, Koboldt DC, Abbott TE, Dooling DJ, et al. SomaticSniper: identification of somatic point mutations in whole genome sequencing data. Bioinformatics. 2012;28:311–7.

19. Saunders CT, Wong WSW, Swamy S, Becq J, Murray LJ, Cheetham RK. Strelka: accurate somatic smallvariant calling from sequenced tumor–normal sample pairs. Bioinformatics. 2012;28:1811–7.

20. Valle ÍF do, Giampieri E, Simonetti G, Padella A, Manfrini M, Ferrari A, et al. Optimized pipeline of MuTect and GATK tools to improve the detection of somatic single nucleotide polymorphisms in whole-exome sequencing data. Bmc Bioinformatics. 2016;17:341.

21. Benjamin DI, Sato T, Lichtenstein L, Stewart C, Getz G, Cibulskis K. Calling Somatic SNVs and Indels with Mutect2. Biorxiv. 2019;861054.

22. Hayward JJ, White ME, Boyle M, Shannon LM, Casal ML, Castelhano MG, et al. Imputation of canine genotype array data using 365 whole-genome sequences improves power of genome-wide association studies. Plos Genet. 2019;15:e1008003.

23. Li H. A statistical framework for SNP calling, mutation discovery, association mapping and population genetical parameter estimation from sequencing data. Bioinformatics. 2011;27:2987–93.

24. Cingolani P, Platts A, Wang LL, Coon M, Nguyen T, Wang L, et al. A program for annotating and predicting the effects of single nucleotide polymorphisms, SnpEff. Fly. 2012;6:80–92.

25. Therneau TM. Survival Analysis [Internet]. 2014. Available from: http://r-forge.r-project.org

26. Kassambara A, Kosinski M, Biecek P. Drawing Survival Curves using “ggplot2” [Internet]. 2019. Available from: http://www.sthda.com/english/rpkgs/survminer/

27. Bista P, Zeng W, Ryan S, Bailly V, Browning JL, Lukashev ME. TRAF3 Controls Activation of the Canonical and Alternative NFκB by the Lymphotoxin Beta Receptor. J Biol Chem. 2010;285:12971–8.

28. Johnson DP, Spitz GS, Tharkar S, Quayle SN, Shearstone JR, Jones S, et al. HDAC1,2 inhibition impairs EZH2- and BBAP-mediated DNA repair to overcome chemoresistance in EZH2 gain-of-function mutant diffuse large B-cell lymphoma. Oncotarget. 2015;6:4863–87.

29. Lawrence MS, Stojanov P, Mermel CH, Robinson JT, Garraway LA, Golub TR, et al. Discovery and saturation analysis of cancer genes across 21 tumour types. Nature. 2014;505:495–501.

30. Akhoondi S, Sun D, Lehr N von der, Apostolidou S, Klotz K, Maljukova A, et al. FBXW7/hCDC4 Is a General Tumor Suppressor in Human Cancer. Cancer Res. 2007;67:9006–12.

31. Kourtis N, Strikoudis A, Aifantis I. Emerging roles for the FBXW7 ubiquitin ligase in leukemia and beyond. Curr Opin Cell Biol. 2015;37:28–34.

32. Ishii N, Araki K, Yokobori T, Gantumur D, Yamanaka T, Altan B, et al. Reduced FBXW7 expression in pancreatic cancer correlates with poor prognosis and chemotherapeutic resistance via accumulation of MCL1. Oncotarget. 2014;5:112636–46.

33. Ye M, Zhang Y, Zhang X, Zhang J, Jing P, Cao L, et al. Targeting FBW7 as a Strategy to Overcome Resistance to Targeted Therapy in Non–Small Cell Lung Cancer. Cancer Res. 2017;77:3527–39.

34. Licht JD. SETD2: a complex role in blood malignancy. Blood. 2017;130:2576–8.

35. Kastenhuber ER, Lowe SW. Putting p53 in Context. Cell. 2017;170:1062–78.

36. Levine AJ. Targeting Therapies for the p53 Protein in Cancer Treatments. Annu Rev Cancer Biology. 2019;3:21–34.

37. Koshino A, Goto-Koshino Y, Setoguchi A, Ohno K, Tsujimoto H. Mutation of p53 Gene and Its Correlation with the Clinical Outcome in Dogs with Lymphoma. J Vet Intern Med. 2016;30:223–9.

38. Taylor-Weiner A, Stewart C, Giordano T, Miller M, Rosenberg M, Macbeth A, et al. DeTiN: overcoming tumor-in-normal contamination. Nat Methods. 2018;15:531–4.

39. Pon JR, Marra MA. Driver and passenger mutations in cancer. Annu Rev Pathology. 2014;10:25–50.

40. Guy MK, Page RL, Jensen WA, Olson PN, Haworth JD, Searfoss EE, et al. The Golden Retriever Lifetime Study: establishing an observational cohort study with translational relevance for human health. Philosophical Transactions Royal Soc B Biological Sci. 2015;370:20140230.

41. Robinson JT, Thorvaldsdóttir H, Winckler W, Guttman M, Lander ES, Getz G, et al. Integrative genomics viewer. Nat Biotechnol. 2011;29:24–6.

