## Supplemental Results for "Consensus-based somatic variant-calling method correlates *FBXW7* mutations with poor prognosis in canine B-cell lymphoma"

### Supplement

#### Results of 4-caller method

In preparation for using a consensus approach of keeping variants called by 2 or more of 4 callers (VarScan 2, Somatic Sniper, Strelka, and MuTect), all callers were run individually. Different calling approaches yielded a six-fold difference in the number of tumor variants identified, with MuTect being the most conservative caller and VarScan 2 being the least. Regardless of which caller, variants identified by only one of the four callers had a high likelihood of being a false positive variant call when aligned reads were manually examined with the Integrated Genomics Viewer (IGV) program (41).

Using these custom consensus parameters for 60 dogs, 9,059 SNPs (151 per dog) were called by 3 or more of the 4 variant callers and 50,210 SNPs (836 per dog) were called by 2 or more variant callers across all 60 tumors. Approximately 800 somatic mutations per dog passing our calling method was consistent with published literature. However, it was unexpected that many of the highest ranked genes (in terms of number of dogs with at least 1 somatic mutation) encoded uncharacterized genes and many of these mutations were silent or in non-coding regions. Visual inspection of calls revealed that a high number of false positive calls were still accepted by our method. Despite an abundance of false positive calls, however, 25% of samples were called as containing at least one exonic mutation in the TRAF3 gene, and in this respect our data was consistent with previous publications.

On average, the 4-caller method kept 836 variants per dog (accepted by 2 or more callers) with an average of 151 per dog called by three or all callers.
