## Supplemental Figure 1 for "Consensus-based somatic variant-calling method correlates *FBXW7* mutations with poor prognosis in canine B-cell lymphoma"

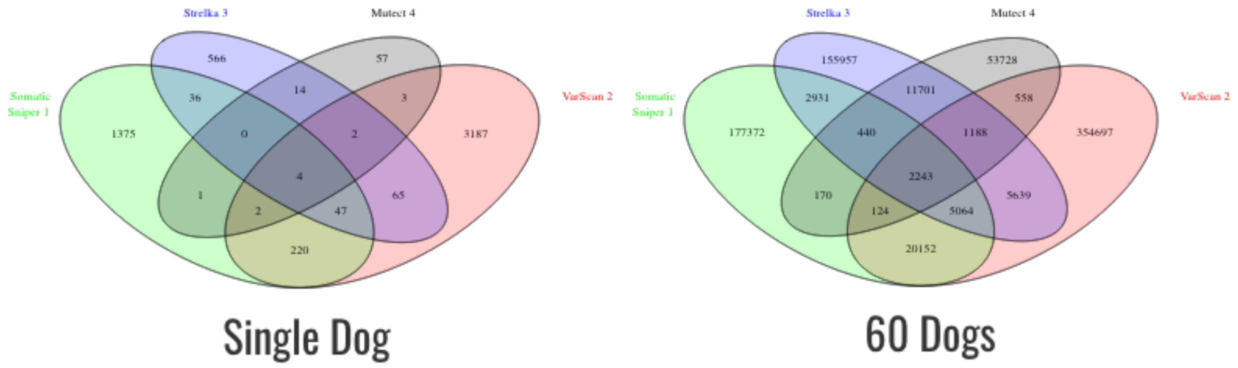


**Supplemental Figure 1: Comparison of number of variants called by single and multiple callers for a single dog and combined for all dogs (60 dogs) using the 4-caller method.**
