## Supplemental Table 1 for "Consensus-based somatic variant-calling method correlates *FBXW7* mutations with poor prognosis in canine B-cell lymphoma"

| Dog ID | Total Mapped Sequences Tumor | Total Mapped Sequences Normal Tissue | 4Caller Mutations | 2Caller Mutations | 4 Caller Variants Per Read | 2 Caller Variants Per Read | Treatment | Achieved Remission? | Treated At Relapse? | PFS | OS | Censored (1 = yes, 2 = no) | | TRAF3 | DIAPH2 | ENSCAFG-00000030258 | POT1 | ENSCAFG-00000029236 | KDM6A | FBXW7 | SETD2 | TP53 |
|---|---|---|---|---|---|---|---|---|---|---|---|---|---|---|---|---|---|---|---|---|---|---|
| 4100 | 40,587,907 | 54,076,022 | 445 | 75 | 1.10E-05 | 1.85E-06 | Multi-agent | Yes | Yes | 1196 | 1354 | 2 | | NO | NO | NO | NO | NO | NO | NO | NO | YES |
| 4101 | 52,126,701 | 41,156,401 | Not Called | 23 | Not Called | 4.41E-07 | Multi-agent | Yes | Yes | 316 | 480 | 2 | | YES | NO | NO | YES | NO | YES | NO | NO | NO |
| 4102 | 50,199,469 | 47,262,808 | 567 | 2 | 1.13E-05 | 3.98E-08 | Single Agent | Yes | Yes | 260 | 260 | 1 | | NO | NO | NO | NO | NO | NO | NO | NO | NO |
| 4103 | 40,706,893 | 32,440,251 | 394 | 9 | 9.68E-06 | 2.21E-07 | Multi-agent | Yes | Yes | 348 | 611 | 2 | | NO | NO | NO | NO | NO | NO | NO | NO | NO |
| 4104 | 50,368,621 | 36,134,585 | 297 | 3 | 5.90E-06 | 5.96E-08 | Induction only | No | NA | NA | 342 | 2 | | NO | NO | NO | NO | NO | NO | NO | NO | NO |
| 4105 | 39,212,233 | 46,540,375 | 370 | 40 | 9.44E-06 | 1.02E-06 | Induction only | No | NA | NA | 4 | 1 | | YES | NO | NO | NO | YES | NO | NO | NO | NO |
| 4107 | 33,811,949 | 46,991,323 | Not Called | 75 | Not Called | 2.22E-06 | Induction only | NA | NA | NA | 249 | 2 | | NO | YES | NO | NO | NO | NO | NO | NO | YES |
| 4108 | 45,708,844 | 30,316,673 | 6727 | 45 | 1.47E-04 | 9.84E-07 | Single Agent | Yes | Yes | 358 | 427 | 2 | | NO | NO | NO | NO | NO | NO | NO | NO | NO |
| 4110 | 32,290,186 | 40,757,682 | 508 | 35 | 1.57E-05 | 1.08E-06 | Multi-agent | Yes | Yes | 159 | 297 | 2 | | YES | NO | NO | YES | NO | NO | YES | YES | NO |
| 4111 | 40,799,781 | 44,340,864 | 411 | 7 | 1.01E-05 | 1.72E-07 | Clinical Data Excluded | | | | | | | NO | NO | NO | NO | NO | NO | NO | NO | NO |
| 4112 | 35,297,848 | 38,784,540 | 6590 | 23 | 1.87E-04 | 6.52E-07 | None | No | NA | NA | 2 | 2 | | NO | NO | NO | NO | NO | NO | NO | NO | NO |
| 4113 | 38,182,282 | 36,608,729 | 371 | 3 | 9.72E-06 | 7.86E-08 | Clinical Data Excluded | | | | | | | NO | NO | NO | NO | NO | NO | NO | NO | NO |
| 4115 | 38,720,300 | 23,383,649 | 924 | 51 | 2.39E-05 | 1.32E-06 | None | No | NA | NA | 3 | 2 | | NO | NO | NO | NO | NO | YES | NO | NO | NO |
| 4116 | 45,346,910 | 57,208,275 | 499 | 64 | 1.10E-05 | 1.41E-06 | Multi-agent | Yes | Yes | 104 | 204 | 2 | | NO | YES | NO | NO | YES | NO | NO | NO | NO |
| 4117 | 47,329,281 | 43,772,172 | 1017 | 56 | 2.15E-05 | 1.18E-06 | Clinical Data Excluded | | | | | | | YES | NO | NO | NO | YES | NO | YES | NO | NO |
| 4118 | 37,579,080 | 67,749,638 | 409 | 50 | 1.09E-05 | 1.33E-06 | Multi-agent | No | NA | NA | 167 | 2 | | NO | NO | YES | NO | NO | NO | NO | NO | YES |
| 4119 | 88,892,916 | 73,312,732 | 1716 | 15 | 1.93E-05 | 1.69E-07 | Single Agent | Yes | None | NA | 584 | 2 | | NO | NO | NO | NO | NO | NO | NO | NO | NO |
| 4120 | 80,031,939 | 71,987,792 | 720 | 118 | 9.00E-06 | 1.47E-06 | Multi-agent | Yes | None | 43 | 43 | 2 | | NO | YES | NO | NO | YES | NO | NO | NO | NO |
| 4121 | 42,435,682 | 64,377,817 | 918 | 11 | 2.16E-05 | 2.59E-07 | Single Agent | No | NA | NA | 31 | 2 | | NO | NO | NO | NO | NO | NO | NO | NO | NO |
| 4122 | 58,596,018 | 55,697,029 | 558 | 27 | 9.52E-06 | 4.61E-07 | Clinical Data Excluded | | | | | | | NO | NO | NO | NO | NO | NO | NO | NO | NO |
| 4123 | 19,456,144 | 77,263,076 | 344 | 6 | 1.77E-05 | 3.08E-07 | Multi-agent | No | NA | NA | 182 | 2 | | NO | NO | NO | NO | NO | NO | NO | NO | NO |
| 4125 | 44,580,913 | 63,371,799 | 418 | 50 | 9.38E-06 | 1.12E-06 | Single Agent | No | NA | NA | 28 | 2 | | NO | YES | NO | NO | YES | NO | NO | NO | NO |
| 4126 | 60,096,528 | 51,802,756 | Not Called | 61 | Not Called | 1.02E-06 | Single Agent | No | NA | NA | 69 | 2 | | YES | NO | YES | NO | NO | NO | YES | NO | YES |
| 4127 | 38,716,220 | 76,480,571 | 320 | 40 | 8.27E-06 | 1.03E-06 | Multi-agent | Yes | Yes | 333 | 554 | 2 | | YES | NO | NO | YES | NO | NO | YES | NO | NO |
| 4128 | 45,948,708 | 62,754,932 | 693 | 50 | 1.51E-05 | 1.09E-06 | Multi-agent | Yes | Yes | 285 | 551 | 2 | | YES | NO | NO | NO | YES | NO | NO | YES | NO |
| 4129 | 52,251,095 | 49,559,851 | 578 | 32 | 1.11E-05 | 6.12E-07 | Multi-agent | Ye | None | 188 | 361 | 2 | | YES | NO | YES | YES | NO | NO | NO | NO | NO |
| 4130 | 56,693,407 | 71,186,366 | 605 | 57 | 1.07E-05 | 1.01E-06 | Single Agent | Yes | None | 131 | 131 | 2 | | YES | NO | NO | NO | NO | NO | NO | NO | NO |
| 4131 | 59,006,624 | 62,541,761 | 806 | 80 | 1.37E-05 | 1.36E-06 | Single Agent | Yes | None | NA | 138 | 2 | | YES | NO | YES | NO | NO | YES | NO | NO | NO |
| 4132 | 33,103,347 | 48,785,947 | Not Called | 42 | Not Called | 1.27E-06 | Multi-agent | Yes | Yes | 288 | 433 | 2 | | YES | YES | NO | NO | NO | NO | NO | NO | NO |
| 4136 | 59,747,359 | 49,950,142 | Not Called | 4 | Not Called | 6.69E-08 | Clinical Data Excluded | | | | | | | NO | NO | NO | NO | NO | NO | NO | NO | NO |
| 4140 | 37,389,305 | 39,815,700 | Not Called | 68 | Not Called | 1.82E-06 | Single Agent | Yes | Yes | 76 | 84 | 2 | | NO | NO | NO | NO | NO | NO | YES | NO | NO |
| 4142 | 56,389,413 | 52,371,331 | 783 | 73 | 1.39E-05 | 1.29E-06 | Multi-agent | Yes | Yes | 97 | 97 | 1 | | YES | YES | NO | NO | NO | YES | NO | NO | NO |
| 4144 | 59,742,571 | 53,871,137 | 1342 | 63 | 2.25E-05 | 1.05E-06 | Multi-agent | Yes | Yes | 132 | 188 | 2 | | NO | NO | NO | NO | NO | YES | NO | NO | NO |
| 4145 | 42,695,893 | 61,469,443 | 452 | 9 | 1.06E-05 | 2.11E-07 | Multi-agent | Yes | None | 224 | 224 | 1 | | NO | NO | NO | NO | NO | NO | NO | NO | NO |
| 4146 | 36,602,852 | 24,018,828 | Not Called | 67 | Not Called | 1.83E-06 | Multi-agent | Yes | Yes | 230 | 288 | 2 | | NO | YES | NO | NO | NO | NO | NO | NO | NO |
| 4147 | 16,237,922 | 28,629,161 | 272 | 9 | 1.68E-05 | 5.54E-07 | Multi-agent | Yes | Yes | 363 | 482 | 2 | | NO | NO | NO | NO | YES | NO | NO | NO | NO |
| 4148 | 33,469,023 | 24,472,354 | 450 | 26 | 1.34E-05 | 7.77E-07 | Multi-agent | Yes | Yes | 415 | 958 | 2 | | NO | NO | YES | NO | NO | NO | NO | NO | NO |
| 4150 | 41,333,082 | 23,916,545 | 737 | 26 | 1.78E-05 | 6.29E-07 | Multi-agent | Yes | Yes | 241 | 365 | 2 | | YES | NO | NO | YES | NO | NO | NO | NO | NO |
| 4151 | 17,385,440 | 18,992,426 | 642 | 18 | 3.69E-05 | 1.04E-06 | Induction only | NA | NA | NA | 57 | 2 | | NO | YES | NO | NO | NO | NO | NO | NO | YES |
| 4152 | 28,161,288 | 30,428,640 | 588 | 26 | 2.09E-05 | 9.23E-07 | Single Agent | NA | NA | NA | 163 | 2 | | NO | YES | NO | NO | NO | NO | NO | YES | NO |
| 4154 | 23,909,951 | 22,397,877 | 348 | 0 | 1.46E-05 | 0 | No treatment | No | None | NA | 2 | 2 | | NO | NO | NO | NO | NO | NO | NO | NO | NO |
| 4155 | 23,605,240 | 56,287,593 | Not Called | 4 | Not Called | 1.69E-07 | Single Agent | Yes | NA | NA | 102 | 1 | | NO | NO | NO | NO | NO | NO | NO | NO | NO |
| 4156 | 13,585,144 | 29,175,793 | Not Called | 4 | Not Called | 2.94E-07 | Induction only | NA | NA | NA | 132 | 2 | | NO | NO | NO | NO | NO | NO | NO | NO | NO |
| 4158 | 1,218,902 | 33,610,285 | 52 | 1 | 4.27E-05 | 8.20E-07 | Multi-agent | Yes | NA | NA | 448 | 2 | | NO | NO | NO | NO | NO | NO | NO | NO | NO |
| 4160 | 19,651,976 | 221,073,797 | 212 | 9 | 1.08E-05 | 4.58E-07 | Multi-agent | Yes | None | 259 | 271 | 2 | | NO | NO | NO | NO | NO | NO | NO | NO | NO |
| 4161 | 27,105,873 | 25,070,099 | 341 | 1 | 1.26E-05 | 3.69E-08 | Multi-agent | Yes | Yes | 260 | 484 | 2 | | NO | NO | NO | NO | NO | NO | NO | NO | NO |
| 4162 | 19,929,113 | 21,523,091 | 474 | 25 | 2.38E-05 | 1.25E-06 | Multi-agent | No | NA | NA | 9 | 2 | | NO | NO | YES | YES | NO | NO | YES | NO | NO |
| 4163 | 28,638,154 | 23,171,874 | 515 | 2 | 1.80E-05 | 6.98E-08 | Multi-agent | Yes | Yes | 598 | 960 | 2 | | NO | NO | NO | NO | NO | NO | NO | NO | NO |
| 4164 | 41,360,660 | 19,536,901 | 579 | 12 | 1.40E-05 | 2.90E-07 | Multi-agent | Yes | None | 110 | 110 | 2 | | NO | NO | NO | NO | NO | NO | YES | NO | NO |
| 4166 | 39,837,071 | 23,888,936 | 868 | 35 | 2.18E-05 | 8.79E-07 | Single Agent | Yes | None | NA | 143 | 2 | | YES | NO | NO | YES | NO | NO | NO | NO | NO |
| 4167 | 22,827,076 | 21,382,834 | 774 | 40 | 3.39E-05 | 1.75E-06 | Multi-agent | Yes | Yes | 43 | 143 | 2 | | NO | NO | YES | NO | NO | YES | NO | YES | NO |
| 4168 | 25,145,148 | 29,182,019 | 407 | 27 | 1.62E-05 | 1.07E-06 | Multi-agent | Yes | None | 194 | 194 | 2 | | NO | NO | NO | NO | NO | YES | NO | NO | NO |
| 4169 | 28,771,778 | 29,521,143 | 398 | 33 | 1.38E-05 | 1.15E-06 | Clinical Data Excluded | | | | | | | NO | NO | NO | NO | NO | NO | NO | NO | YES |
| 4170 | 34,124,896 | 37,061,449 | 547 | 11 | 1.60E-05 | 3.22E-07 | Clinical Data Excluded | | | | | | | NO | NO | NO | NO | NO | NO | NO | NO | NO |
| 4171 | 51,226,600 | 34,130,517 | 854 | 29 | 1.67E-05 | 5.66E-07 | Clinical Data Excluded | | | | | | | NO | NO | YES | NO | NO | NO | NO | NO | NO |
| 4172 | 21,813,310 | 44,777,015 | 659 | 31 | 3.02E-05 | 1.42E-06 | Clinical Data Excluded | | | | | | | NO | NO | NO | NO | NO | NO | NO | YES | NO |
| 4173 | 54,487,229 | 23,842,748 | 806 | 12 | 1.48E-05 | 2.20E-07 | Multi-agent | No | NA | NA | 94 | 2 | | NO | NO | NO | NO | NO | NO | NO | NO | NO |
| 4174 | 54,209,250 | 64,644,380 | 707 | 39 | 1.30E-05 | 7.19E-07 | Multi-agent | No | NA | NA | 54 | 2 | | NO | NO | NO | NO | NO | NO | YES | YES | NO |
| 4175 | 44,135,308 | 43,021,542 | 555 | 46 | 1.26E-05 | 1.04E-06 | Single Agent | Yes | Yes | 55 | 111 | 2 | | NO | NO | NO | NO | YES | NO | NO | NO | NO |
| 4178 | 17,906,874 | 45,112,091 | 278 | 38 | 1.55E-05 | 2.12E-06 | Multi-agent | Yes | None | 837 | 837 | 2 | | NO | NO | NO | NO | YES | YES | NO | YES | NO |
| 4179 | 32,479,154 | 48,708,307 | 645 | 37 | 1.99E-05 | 1.14E-06 | Single Agent | NA | NA | NA | 62 | 2 | | YES | NO | NO | NO | NO | NO | NO | NO | NO |
| 4181 | 37,733,382 | 61,705,423 | 422 | 59 | 1.12E-05 | 1.56E-06 | Multi-agent | Yes | Yes | 162 | 169 | 2 | | YES | YES | NO | NO | NO | NO | NO | NO | NO |
| 4184 | 43,363,621 | 113,492,179 | 768 | 61 | 1.77E-05 | 1.41E-06 | Multi-agent | Yes | Yes | 283 | 384 | 2 | | YES | NO | YES | YES | NO | NO | NO | YES | NO |
| 4186 | 17,526,697 | 36,298,509 | 309 | 91 | 1.76E-05 | 5.19E-06 | Multi-agent | Yes | NA | 1603 | 1613 | 1 | | NO | YES | NO | YES | NO | NO | NO | NO | NO |
| 4188 | 34,457,026 | 96,477,517 | 939 | 44 | 2.73E-05 | 1.28E-06 | Multi-agent | Yes | None | 80 | 225 | 2 | | YES | NO | YES | NO | NO | NO | NO | NO | NO |
| 4189 | 15,317,857 | 66,314,548 | 2559 | 126 | 1.67E-04 | 8.23E-06 | Multi-agent | No | Yes | NA | 109 | 2 | | NO | NO | NO | NO | NO | NO | NO | NO | NO |
| 4191 | 5,561,714 | 46,520,663 | 185 | 4 | 3.33E-05 | 7.19E-07 | Single Agent | No | NA | NA | 77 | 1 | | NO | NO | NO | NO | NO | NO | NO | NO | NO |
| 4194 | 55,938,991 | 40,783,797 | 688 | 43 | 1.23E-05 | 7.69E-07 | Single Agent | No | NA | NA | 38 | 2 | | NO | YES | NO | NO | YES | NO | NO | NO | NO |
| 4195 | 43,946,345 | 43,247,404 | 905 | 29 | 2.06E-05 | 6.60E-07 | Multi-agent | No | NA | NA | 196 | 1 | | YES | NO | NO | NO | NO | YES | NO | NO | NO |
| 4197 | 64,577,297 | 48,631,245 | 478 | 16 | 7.40E-06 | 2.48E-07 | Multi-agent | Yes | 1 | 91 | 147 | 2 | | NO | NO | NO | NO | NO | NO | YES | NO | NO |
| 4199 | 61,180,636 | 57,107,157 | 467 | 42 | 7.63E-06 | 6.86E-07 | Multi-agent | Yes | NA | 1122 | 1141 | 2 | | YES | NO | YES | YES | NO | NO | NO | NO | NO |
| **AVERAGE** | **39,194,426** | **48,341,984** | **810** | **35** | **2.36E-05** | **1.01E-06** | | | | **323** | **292** | | **Total** | **20** | **12** | **11** | **10** | **10** | **9** | **9** | **8** | **6** |

**Supplementary Table 1: Variant calls and Clinical Data by Dog.**

Select genes listed with mutation status (yes/no for presence/absence of a called mutation).
